# A mouse model of hemochromatosis-related mutations with brain iron dyshomeostasis exhibits loss of tyrosine hydroxylase expression in dopaminergic neurons and motor control impairment relevant to Parkinson’s disease

**DOI:** 10.1101/2025.05.06.648286

**Authors:** Elvis Freeman-Acquah, Rebecca Hood, Chan-An Lin, Qiao-Xin Li, Luke Gordon, Kristy Martin, Ritambhara Aryal, Adrienne E. Milward, Daniel M. Johnstone

**Author notes:** **Corresponding Authors:** Daniel Johnstone, PhD, Adrienne Milward, PhD.

## Abstract

UK Biobank studies show Parkinson’s disease risk is almost doubled in men homozygous for the homeostatic iron regulator gene *HFE* p.C282Y polymorphism, associated with the common genetic iron disorder hemochromatosis. Whether this relationship is causal or spurious is unknown. We previously reported a novel *Hfe*^*-/-*^*xTfr2*^*mut*^ mouse model of hemochromatosis with elevated brain iron (∼1.5-1.8x). We now show these mice have reduced substantia nigra tyrosine hydroxylase expression at 3 months and 9 months age, sometimes exhibit severe hindlimb clasping by 7-8 months, have impaired rotarod and balance beam performance at 9 months and are untestable on the pole test. These parkinsonian features place the model at the forefront of genetic mouse models of PD, which generally do not show both TH loss and motor impairment. This confirms hemochromatosis-related mutations can cause parkinsonian features, substantiating causality of epidemiological relationships. Despite total brain iron elevation, neuronal iron remains low in *Hfe*^*-/-*^*xTfr2*^*mut*^ mice, consistent with hemochromatosis-related mutations disrupting the normal, iron-responsive regulation of the neuronal iron exporter ferroportin by hepcidin. Parkinsonian features may reflect reduced mitochondrial respiratory complex (MRC) activity due to functional neuronal iron depletion. This may be exacerbated by indiscriminate chelation and could instead respond to drugs targeting the hepcidin-ferroportin axis or MRC activity. This new model of chronic parkinsonism that increases with age provides unprecedented insights into the complex relationships of brain iron regulation and movement impairment. Since parkinsonism of diverse etiologies can exhibit iron dysregulation, the model may facilitate pre-clinical to end-stage studies relevant to both sporadic and genetic PD.

## 1 Introduction

The p.C282Y variant in the homeostatic iron regulator protein HFE, carried by ∼10-15% of people of Anglo/central European background, causes over 90% of cases of the common iron disorder hemochromatosis [1,2]. Recently, this same variant has emerged as a strong genetic risk factor for late-life movement impairment and Parkinson’s disease (PD) in men. Epidemiological analysis of large UK BioBank community cohorts (n>335,000, age 49-87y) has provided evidence that men homozygous for HFE p.C282Y polymorphism have almost twice the risks of PD or other movement impairment compared to men without this polymorphism (OR 1.80; 95% CI, 1.28-2.55; p=.001; n=223,569 men, including 1320 p.C282Y homozygotes) [3]. Neuroimaging of a sub-cohort of 165 p.C282Y homozygotes (66 men) and 671 matched controls (272 men) showed homozygosity was associated with decreased T2-weighted and T2* signal intensity in subcortical motor structures (e.g. basal ganglia, thalamus, cerebellum), consistent with substantial iron deposition [3]. An independent UK BioBank analysis of PD alone also found increased risk among p.C282Y homozygous men, with cumulative incidence of 3.7% vs 2.4% in men without mutations (HR 1.86, 95% CI: 1.21 to 2.87 p=0.005; n=133,771; 1298 p.C282Y homozygotes) [4]. This was independent of hemochromatosis diagnosis, leading to calls for pre-emptive iron chelation in p.C282Y homozygous men. Menstruation may help protect women against iron accumulation but risks for women prematurely amenorrheic due to surgical procedures or other causes are unknown.

These new epidemiological data support case reports and clinical studies of movement disorders in hemochromatosis patients [5-7] and mitigation of certain signs (e.g. cerebellar ataxia) in HFE hemochromatosis following iron removal (venesection) [8]. An increased risk for Parkinson’s disease (PD) has also been demonstrated in a case series of patients with hemochromatosis [9].

Yet a recent clinical trial of the iron chelator deferiprone in early PD found that chelation was associated with worse outcomes than placebo control, despite reducing nigrostriatal iron, with dopaminergic therapy initiated due to symptom progression in 22.0% vs. 2.7% of participants [10]. This shows iron chelation monotherapy can adversely affect brain function in PD patients with normal body iron status, most of whom will not be HFE p.C282Y homozygous. People with inherent deficits in iron regulatory mechanisms due to HFE p.C282Y homozygosity may respond differently to chelation therapy than people without HFE mutations, particularly those with elevated body or brain iron levels. Studies in experimental models are needed to assess whether chelation benefits outweigh negative effects in such individuals to determine whether further clinical trials are warranted for some or all HFE pC282Y homozygous individuals. It is also important to test whether alternative treatments may restore normal iron status more safely in these people.

One way to potentially circumvent these issues would be to use the *Hfe*^*-/-*^*xTfr2*^*mut*^ model of chronic and progressive iron overload that our group has created. The model mimics the relatively common human condition of hereditary hemochromatosis. Further, brain iron loading in this model results from the systemic phenomenon of excessive dietary iron absorption, rather than from localized mishandling, as a result of rare faults in iron-related proteins. Powerful experimental models are needed to understand the complexities of brain iron homeostasis and relationships with disease risks in genetic hemochromatosis.

We have performed foundational studies in our unique genetic mouse model of brain iron dyshomeostasis caused by hemochromatosis-related mutations [11,12] offering fresh insights into this question and unprecedented opportunities to test new drugs directed at hemochromatosis-related targets. In this study we use a combination of chronic, progressive iron overload in *Hfe*^*-/-*^*xTfr2*^*mu*t^ mouse model of hemochromatosis to determine effects of hemochromatosis-related mutations relevant to PD.

## 2 Materials and methods

The data that support the study findings are available from the corresponding author upon reasonable request.

### 2.1 Animals and ethical statement

This study used the *Hfe*^*-/-*^ *xTfr2* ^*mut*^ mouse model of brain iron elevation (‘Iron’ model) on an AKR/J background and strain-matched wildtype controls. Since UK Biobank data only showed effects in males the following studies were conducted on male mice only. All protocols were approved by the Animal Ethics Committee of the University of Sydney (2017/1188) and followed the guidelines of the National Health and Medical Research Council, Australia. Procedures were performed on mice reared at the University of Sydney animal facility. All mice were housed in individually ventilated cages at 22-24°C ambient temperature with *ad libitum* access to both water and food.

### 2.2 Experimental design

#### 2.2.1 Study 1: Testing effects of hemochromatosis-related mutations on movement in male mice

To determine whether hemochromatosis-related mutations have sex specific effects on motor function, *Hfe*^*-/-*^*xTfr2*^*mu*t^ (n=6) and wildtype controls (n=5-6) were screened at 9 months of age for PD-like features by assessing balance and motor coordination using the rotarod and balance beam tasks (Sections 2.3.1, 2.3.2 below).

#### 2.2.2 Study 2: Effects of hemochromatosis-related mutations on brain iron homeostasis

Histochemical labeling with 3, 3’ diaminobenzidine tetrahydrochloride (DAB)-enhanced Perls’ stain was used to detect ferric (Fe3+) iron in substantia nigra pars compacta (SNc) in brain cryosections from mice at 9 months of age (wildtype n=5; *Hfe*^*-/-*^*xTfr2*^*mu*t^ n=5).

#### 2.2.3 Study 3: Effects of hemochromatosis-related mutations on movement control

To replicate and further characterize effects of hemochromatosis-related mutations on movement control, 3 month and 9 month old wildtype and *Hfe*^*-/-*^*xTfr2*^*mu*t^ mice (3 months: wildtype n=6; *Hfe*^*-/-*^ *xTfr2*^*mu*t^ n=5; 9 months: wildtype n=5; *Hfe*^*-/-*^*xTfr2*^*mu*t^ n=5).

The age of 3 months was selected to represent bona fide, fully mature adult mice, in accordance with Jackson Laboratory recommendations [13], and because we previously observed brain iron is increased ∼1.5x in *Hfe*^*-/-*^*xTfr2*^*mu*t^ mice relative to wildtype controls at this age [11,12,14], minimizing the possibility of false negative outcomes at this age due to confounding ceiling effects.

#### 2.2.4 Study 4: Effects of hemochromatosis-related mutations on tyrosine hydroxylase (TH) expression in SNc neurons

To determine the effect of hemochromatosis-related mutations on the nigrostriatal pathway, loss of dopaminergic cells and their terminals was assessed at both ages by immunolabeling of the SNc with the dopaminergic cell marker tyrosine hydroxylase (TH). The age of 3 months was selected to represent bona fide, fully mature adult mice in accordance with Jackson Laboratory recommendations [13]; we have previously shown iron an attempt to avoid any confounding ceiling effects that may occur in the older mice.

#### 2.2.5 Study 5: Alpha Synuclein

Western immunoblotting was used to determine the effects of hemochromatosis-related mutations on levels of *α*-synuclein in brain homogenates from mice age 9 months.

### 2.3 Motor Assessment

The rotarod test [15-19] and balance beam [17,20] were used to assess balance and motor coordination.

#### 2.3.1 Rotarod

Assessment of balance and motor coordination using the rotarod was performed as previously described [21]. Briefly, mice were acclimatized to the room for 30 min, followed by habituation to the rotarod for 2 min, before being returned to their cage for 5 min. During testing, the mice were placed on the static cylindrical rods and the trial commenced at a starting speed of 5 rpm. Rotation speed accelerated to 20 rpm over 120 seconds, at which time the trial ended. Each of the mice performed 5 trials, with 5 min of resting time between each trial. The primary outcome measure was the latency to fall from the rotating rod. Activity was recorded using a video camera for manual analysis.

#### 2.3.2 Balance beam

Assessment of balance and motor coordination using the balance beam was done as previously described [20]. Briefly, on the day prior to testing, mice were habituated and trained to walk on the wooden balance beam (12 mm in diameter, 60 cm above ground level, walking distance 60 cm). During testing, mice were placed at the start of the beam and their first continuous movement (without falling) along the beam for each trial was recorded. Each mouse underwent 5 trials, with an inter-trial interval of 10 min. Activity was recorded using a video camera and videos were analyzed manually. Outcome measures were the time to traverse.

### 2.4 Tissue processing

Mice were euthanized at 3 or 9 months of age by intraperitoneal injection of 60 mg/kg sodium pentobarbitone (Birvac Australia Pty Ltd). Mice were transcardially perfused either with saline, followed by brain collection and snap freezing in liquid nitrogen before storage at -80°C, or with 4% paraformaldehyde, followed by brain collection and storage in 4% paraformaldehyde solution at 4°C for 24 hr. Fixed brain tissue was then transferred into 0.1% sodium azide in 0.1M phosphate buffered saline (PBS) (pH 7.4) and stored at 4°C.

Fixed brains were then bisected along the midline and the left hemisphere assigned to histological examination, while the right hemisphere was stored in the PBS-azide solution for future use. The left hemispheres were cryoprotected with 30% sucrose in 0.1M PBS (pH 7.4), for at least 48 hr or until sucrose had fully infiltrated the tissue. Before embedding, hemispheres were sliced in the coronal plane at the inferior colliculus (-5.00 mm Bregma) and at -2.25 mm Bregma to produce a tissue block containing the SNc (-4.75 mm to -2.52 mm Bregma).

Tissue blocks were embedded in TissueTek (O.C.T compound, SAKURA) and coronally sectioned at 25 μm thick using a cryostat (Leica CM3050; Leica Biosystems). Sections were collected on onto slides coated with poly-L-lysine hydrobromide as a 1:3 series.

### 2.5 Tissue staining and IHC

#### 2.5.1 DAB-enhanced Perls’ staining and visualization of iron

A subset of sections were prepared for DAB-enhanced Perls’ (DAB+Perls’) staining of ferric iron (Fe3+) as previously described [22]. Briefly, slide-mounted SNc sections were removed from PBS and transferred into 1% potassium ferrocyanide (pH <1) and incubated at room temperature for 30 min, before 3×2 min washes in ddH_2_O. Sections were then transferred into a solution of 0.01 M sodium azide and 0.3% hydrogen peroxide in methanol and incubated at room temperature for 1 hr, before 3×2 min washes in PBS. Sections were next incubated in 0.025% DAB (MP Biomedical) and 0.005% hydrogen peroxide in PBS at room temperature for 30 min and then 3×2 min washes in ddH_2_O. After staining, the sections were dehydrated using a graded series of EtOH (50%, 70%, 90%, 2x100%; 2 min each) and cleared in two 5 min exchanges of 100% xylene. Coverslips were applied and allowed to dry overnight before microscopy. Sections were visualized and scanned using the Axioscan (Zeiss) at 20x magnification.

#### 2.5.2 Immunohistochemical analysis of the dopaminergic neuron marker tyrosine hydroxylase

Sections were prepared for IHC as previously described [23]. Briefly, slides were dried at 37°C for 1 hr, before 5 min immersion in 70% EtOH to remove TissueTek from the sections followed by agitation in 1% Triton X-100 (TX-100) in PBS for 2 min in a humidified container. To block non-specific binding, excess TX-100 was tapped off and sections incubated in 10% normal goat serum for 1 hr at RT on a shaker then immersed in 1% Triton X-100 and agitated for 1 hr. Excess TX-100 was tapped off and sections incubated in rabbit anti-tyrosine hydroxylase (TH) primary antibody (1:500; T8700-1VL, Sigma-Aldrich USA, ∼ 30 μl /section) 1 hr at RT on a shaker, followed by incubation at 4°C for 48 hr.

Slides were then washed for 5 min in PBS and incubated in anti-rabbit biotinylated secondary antibody (Vectastain, Elite ABC Kit, Vector Laboratories USA, ∼30 μl/section) for 4 hr at room temperature on a shaker. After secondary antibody incubation, slides were incubated in streptavidin peroxidase tertiary antibody (Vectastain, Elite ABC Kit, Vector Laboratories USA, ∼30 μl/section) for 2 hr at room temperature on a shaker. After incubation, slides were washed for 2 min in PBS, followed by immersion in 0.05% DAB in PBS for 10 min. Hydrogen peroxide was then added to the DAB solution at a final concentration of 0.0001% and slides incubated in this solution 10 min. Slides were then washed in PBS for 2 min, dehydrated by immersing in a graded series of EtOH (50, 70%, 90%, 2x100%; 30 seconds each), cleared using two exchanges of 100% histolene (5 min per exchange), coverslips mounted using DPX (BDH Chemicals) and allowed to dry at RT overnight. Visualization and scanning of the brightfield brown colored iron deposits on DAB+Perls’ stained brain sections were obtained using the Axioscan (ZEISS AxioScan Z.1) at 40x magnification.

#### 2.5.3 Quantification of TH-positive cells in the SNc

The number of TH-positive (TH^+^) cells in the SNc was estimated by stereology using the optical fractionator method (StereoInvestigator, MBF Science). A 1:3 series of sections spanning the entire volume of the SNc (from Bregma -3.88 mm to Bregma -2.54 mm; 11-13 sections each separated by 75 μm) was assessed using an unbiased protocol based on the optical fractionator and optical disector principles [24,25]. Standardized predetermined intervals [(counting frame or optical disector size of x = 45 μm and y = 108 μm and systematic random sampling (SRS) grid of x = 140 μm y = 140 μm)] were kept consistent for all counts. The criteria for cells to be counted were clear TH labeling and visible cell bodies with distinct nucleus within each optical disector at a magnification of 40x. After each count, an estimated population of TH^+^ cells generated by the optical fractionator was recorded and saved.

### 2.6 Western immunoblotting

#### 2.6.1 Sample preparation for biochemical analyses

Whole brain tissue was ground in a mortar pre-cooled with liquid nitrogen and diluted to a 10% homogenate (100 mg/mL) in radioimmunoprecipitation assay buffer (RIPA; 10 mM Tris pH 7.4, 150 mM NaCl, 10 mM EDTA, 0.5% Nonidet P-40, 0.5% sodium deoxycholate). Protein concentration was measured in a NanoDrop One spectrophotometer (Thermo Fisher Scientific). Homogenates were sonicated, centrifuged, and aliquots of supernatant (100μg protein/sample) made up to 50 μL with 1x working sample buffer comprised of 12.5 μL of 4x sample buffer containing Coomassie G250 blue dye and phenol red (both Invitrogen), 32.5 μL ddH2O and 5 μL 500 mM dithiothreitol (Invitrogen) and centrifuged briefly to ensure mixing. Aliquoted samples were stored at 4°C until running of gels.

#### 2.6.2 Electrophoresis and Western transfer

Samples were heated at 90°C for 5 min, microfuged 5 seconds before gel loading and electrophoresed on 4-12% tricine gradient gels (Novex) in 1x MES running buffer (50mM MES, 50mM Tris, 0.1% SDS, 0.03% EDTA). Gels were loaded with sample (20μg/lane), molecular weight markers (SeeBlue Plus2 LC5925, Invitrogen) and sample buffer as negative control and run at 120V. Gels were transferred onto nitrocellulose membrane in ice-cold transfer buffer (25 mM Tris, 192 mM glycine) at 120V for ∼30 min with continuous stirring. After transfer, the membrane was microwaved in 1x PBS for 30 seconds.

#### 2.6.3 Immunolabeling and imaging

The membrane was then stained for total protein with 3% Ponceau S, imaged, rinsed and transferred to 0.5% casein in ddH2O blocking buffer (Sigma-Aldrich) and incubated with rocking for 30 mins, RT. Casein solution was discarded and membrane rinsed briefly with ddH2O. The membrane was incubated overnight at 4°C with rocking in primary antibody (rabbit polyclonal antibody 97/8), as previously described [26]. Next day, the membrane was washed with PBST for 1 cycle (3 x 5 min) and incubated for 1 hr with rocking in secondary antibody (anti-rabbit horse radish peroxidase conjugate, 1:10,000) followed by another washing cycle. Blots were developed for 1 min with a 1:1 mixture of GE Health Prime Chemiluminescence Reagent and SuperSignal West Dura Extended Duration Substrate (ThermoFisher Scientific), then imaged on the Fuji LAS-4000.

#### 2.6.4 Immunoblot analysis

Western blots images were imported into ImageJ and the synuclein and actin bands outlined in each lane using the rectangle selection tool. The average signal value of the band in the lane was measured along with a size-matched background area. Standardization for loading variation was performed by dividing the background corrected *α*-synuclein signal values by the background corrected actin value in the same lane.

### 2.7 Statistical analysis

Statistical analysis and data plotting were performed in GraphPad Prism 10 (GraphPad Software). Comparisons between genotype groups used two-tailed unpaired Student’s t-tests. Two-way ANOVA was used to assess the effect of genotype and age, with Fischer’s LSD test used for post-hoc pairwise comparisons. All data are presented as mean ± SEM.

## 3. Results

### 3.1 Effect of hemochromatosis-related mutations on motor coordination and balance

To determine whether hemochromatosis-related mutations have effects on motor function, we compared rotarod and balance beam performance in male AKR/J wildtype and strain-matched *Hfe*^*-/-*^*xTfr2*^*mut*^ mice at 9 months of age. This age was chosen as the timepoint likely to have greatest iron loading without significant mouse attrition, as our earlier published and preliminary observations suggest that: i. death in our AKR/J background strain, with or without hemochromatosis-related mutations, generally occurs at ∼10-12 months of age on, with few mice surviving to ages 13-15 months (unpublished) though most survive beyond the median survival of ∼8-9.5 months and upper 95% CI range of 10.7 months reported for this strain by the Jackson Aging Laboratory [13]; ii. brain iron, assessed by diverse methods (non-heme iron assay, histochemistry, inductively coupled atomic emission spectroscopy, ferritin immunoblotting, synchrotron X-ray fluorescence and X-ray absorption near edge structure analyses) was greater for wildtype and *Hfe*^*-/-*^*xTfr2*^*mu*t^ mice at 9 months compared to 3 months (e.g. [11,12,14]) or 6 months [27], with *Hfe*^*-/-*^*xTfr2*^*mu*t^ mice greater than wildtype at all ages examined (Fig. 3.7; [14]). iii. some though not all *Hfe*^*-/-*^*xTfr2*^*mu*t^ mice showed severe hindlimb clasping [28] by age 7-8 months, with full retraction of both hindlimbs for most of the duration of suspension (Fig. 1A, B and Suppl. Videos 1 and 2), and *Hfe*^*-/-*^*xTfr2*^*mu*t^ mice were also untestable by the pole test, being unable to orient downward and remain on the pole, consistent with movement impairment on par with or greater than some reported 6-hydroxydopamine (6-OHDA) or 1-methyl-4-phenyl-1,2,3,6-tetrahydropyridine (MPTP) neurotoxin models of striatal dopaminergic depletion [29].

**Figure 1.**
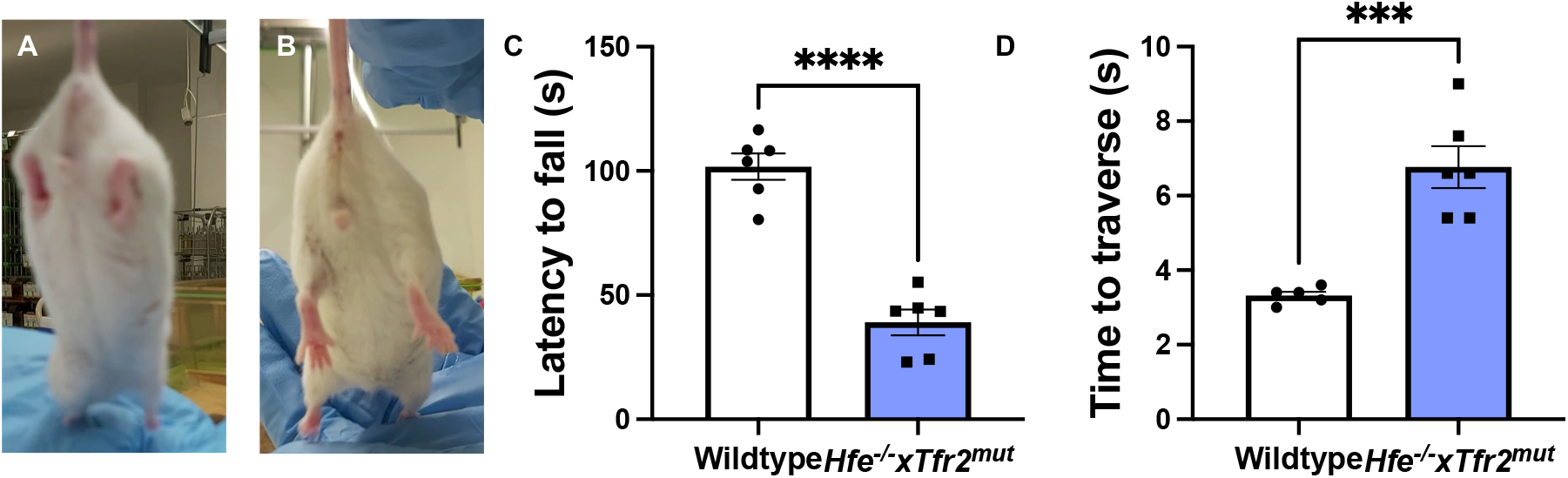
Effect of hemochromatosis-related mutations on motor coordination and balance in male mice. Normal hindlimb splay **(A)** is absent from *Hfe*^*-/-*^*xTfr2*^*mut*^ mice by 7-8 months **(B)**. Motor coordination and balance were assessed by **(C)** duration of time spent on the rotarod, **(D)** the average time spent in traversing the balance beam. Group summary data are presented for both Wildtype and *Hfe*^*-/-*^*xTfr2*^*mut*^ mice at 9 mo. Data are presented as mean ± SEM, *n* = 5-6 mice per group. \*\*\**p*<0.0005, \*\*\*\**p*<0.0001.

#### 3.1.1 Rotarod

Student’s t-test revealed that *Hfe*^*-/-*^*xTfr2*^*mut*^ mice had a significantly shorter average latency to fall compared with wildtype counterparts (*p*<0.0001), with wildtype mice remaining on the rotarod for more than twice the time of Iron mice (Fig. 1C).

#### 3.1.2 Balance Beam

Student’s t-test revealed that *Hfe*^*-/-*^*xTfr2*^*mut*^ mice took twice as long to traverse the beam as wildtype controls (*p*<0.0005; Fig. 1D).

### 3.2 Iron detection by DAB-enhanced Perls’ staining

As previously reported, *Hfe*^*-/-*^*xTfr2*^*mut*^ mice have increased iron labeling compared to control wildtype mice [11,12]. Fig. 2 depicts the distribution and localization of iron detected by DAB+Perls’ staining of coronal brain sections including the SNc from 9-month old male wildtype and *Hfe*^*-/-*^*xTfr2*^*mut*^ mice.

**Figure 2.**
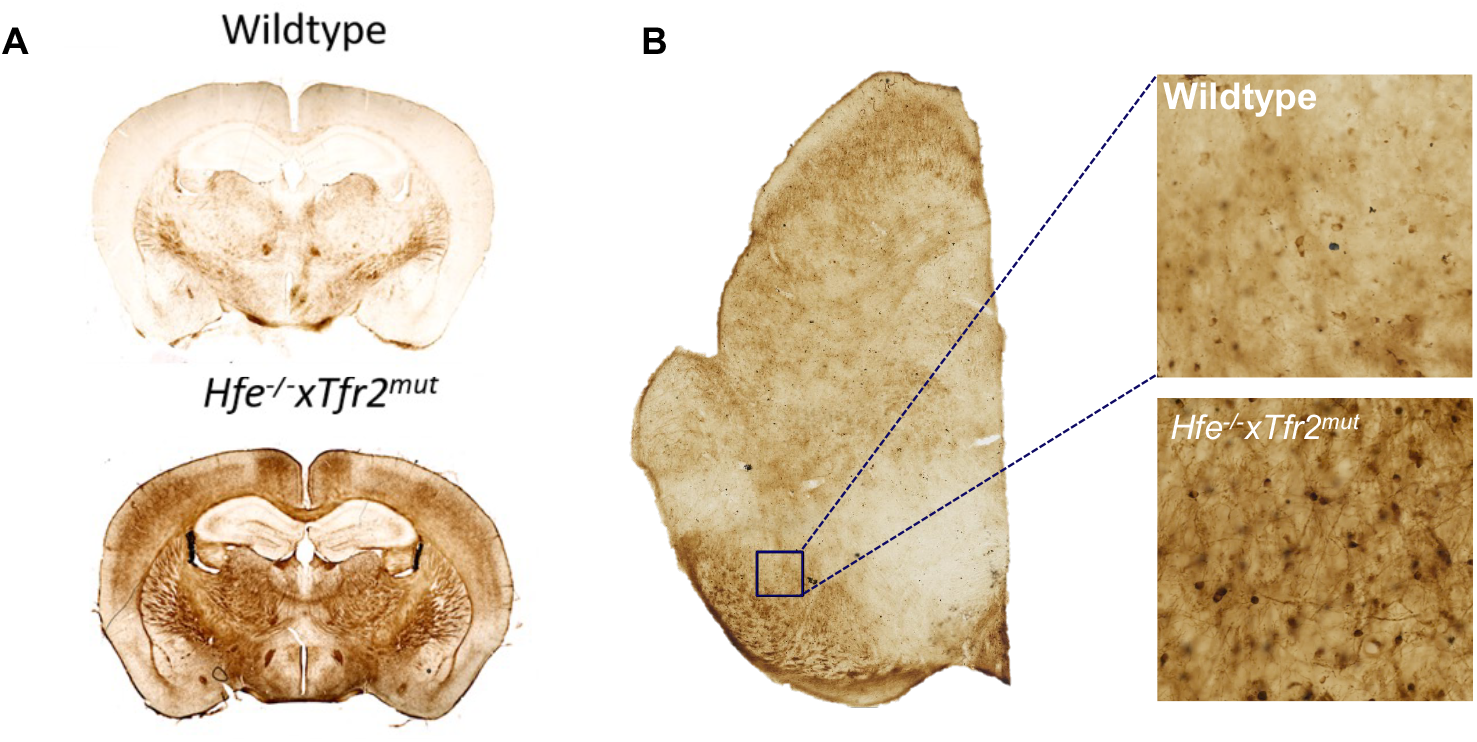
Detection of brain iron by DAB+Perls’ stain in 9-month old male mice. Iron deposition in **(A)** coronal sections of wildtype vs *Hfe*^*-/-*^*xTfr2*^*mut*^ mice and **(B)** the substantia nigra pars compacta (SNc) as determined by 3, 3-diaminobenzidine tetrahydrochloride (DAB)+Perls’ staining.

### 3.3 Dopaminergic cell counts in the substantia nigra pars compacta

Two-way ANOVA revealed a significant effect of genotype (F_(1,17)_=16.73, *p*=0.0008) and age (F_(1,17)_=7.306, *p*=0.0151) on dopaminergic cell count in the SNc. The number of TH^+^ cells was reduced in the SNc of male mice with hemochromatosis-related mutations compared to wildtype mice at both 3 months (2489±192 vs. 3177±128, *p*=0.0073) and 9 months (2716±163 vs. 2069±173, *p*=0.0139; Fig. 3). Post-hoc testing did not show any differences between the 3 and 9 month groups (*p*>0.05).

**Figure 3.**
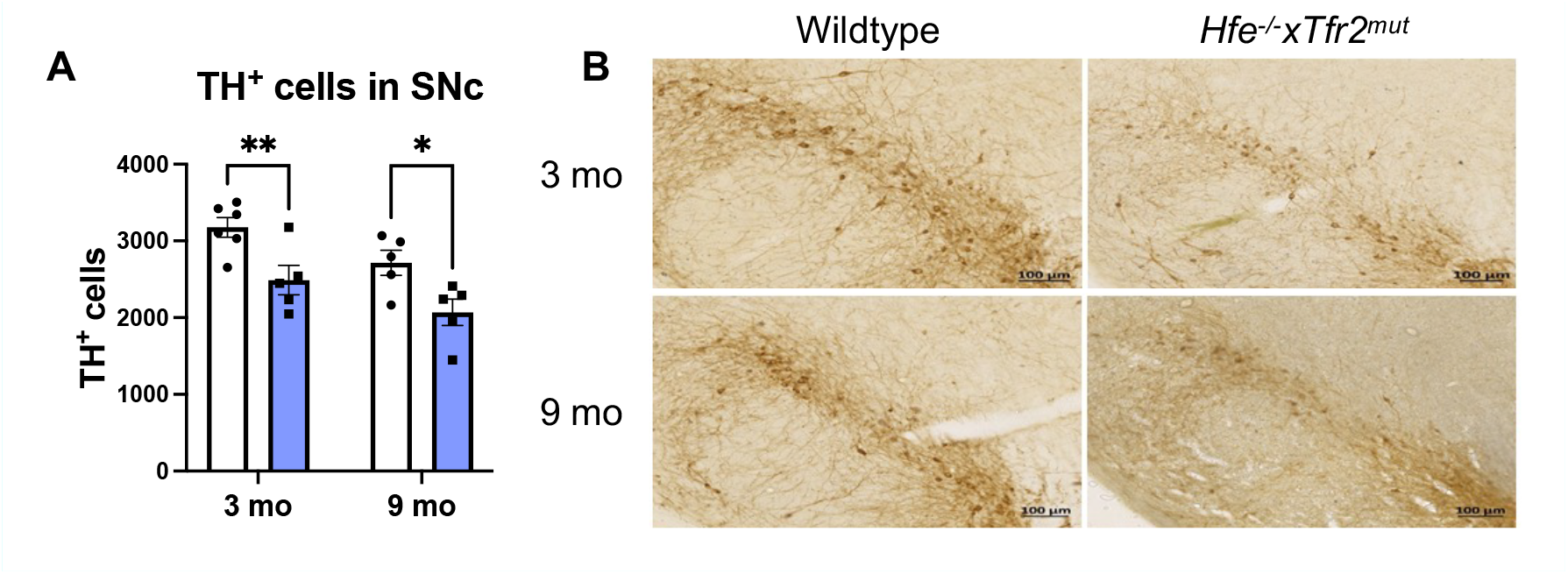
Effects of hemochromatosis-related mutations on TH^+^ cell numbers in the SNc of male mice. **(A)** Stereology was used to assess the number of TH-positive cells in the SNc. Group summary data of TH-positive cell counts are presented for mice aged 3 or 9 months. Data are presented as mean ± SEM, *n* = 5-6 mice per group. \**p*<0.05, \*\**p*<0.01. **(B)** Representative micrographs of TH-labeling of the SNc at 3 and 9 months. Scale bar = 100 µm

### 3.4 Effects of hemochromatosis-related mutations on brain *α*-synuclein levels in male mice

As shown in Fig. 4, mice with hemochromatosis-related mutations had 47% higher levels of *α*-synuclein (1.391±0.119, n=5) compared with wildtypes (0.944±0.053, n=8; *p*=0.0165).

**Figure 4.**
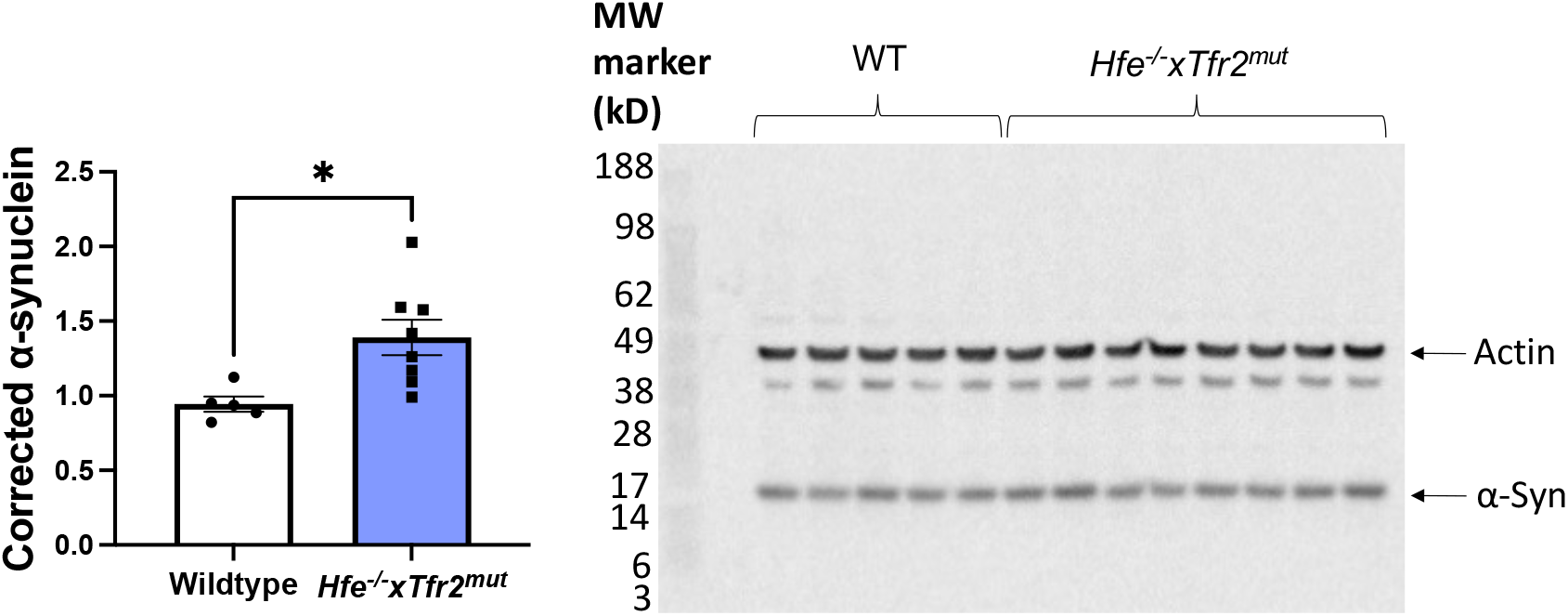
Effect of hemochromatosis-related mutations on α-synuclein levels in male mouse brain. **(A)** Standardized levels of α-synuclein detected by rabbit polyclonal antibody 97/8 to the human α-synuclein C-terminal region 116-131 [26], relative to actin, for homogenized brain samples from *Hfe*^*-/-*^*xTfr2*^*mut*^ mice compared to wildtype mice. **(B)** Western immunoblot visualized by electrochemiluminescence.

## 4 Discussion

The experiments described here explored whether hemochromatosis-related mutations influence behavioral and neuropathological features associated with parkinsonism. Compared to age- and strain-matched male wildtype mice *Hfe*^*-/-*^*xTfr2*^*mut*^ mice had reductions in SNc dopaminergic TH^+^ neurons at both 3 months and 9 months of age, increased brain levels of *α*-synuclein and substantial motor function deficits at 9 months of age. To the best of our knowledge, this study of the *Hfe*^*-/-*^*xTfr2*^*mu*t^ mouse model is the first research to combine assessment of dopaminergic neurons with complementary motor function assessment in a mouse model of hemochromatosis and the first to use a model of hemochromatosis or brain iron elevation *per se* that exhibits chronic, progress brain iron elevation, confirmed directly as opposed to using surrogate measures such as ferritin [30-33].

Hemochromatosis is a common genetic disorder compared to conditions caused by the rare mutations previously shown to cause parkinsonian features that form the basis for the strongest existing mouse genetic models of PD. Furthermore, as far as we know, no other reported genetic mouse model exhibits parkinsonian features across all three domains of loss of dopaminergic TH+ neurons, motor impairment and *α*-synuclein elevation [34,35], with the possible exception of phospholipase A2 *Pla2g6* (PARK14) depletion models of PLA2G6-related dystonia-parkinsonism, from the ‘Neurodegeneration with Brain Iron Accumulation’ (NBIA) rare neurogenetic disease family. We earlier reported decreases in *Pla2g6* transcripts (1.6-fold, *p*<0.01) and altered brain expression of several other NBIA genes in the *Hfe*^*-/-*^ *xTfr2*^*mut*^ model [12].

The findings are consistent with a previous study showing motor function impairment, including rotarod performance, in 3-month old mice with the *Hfe*^*-/-*^ hemochromatosis-related mutation on C57BL6 background [36]. However, while manifesting increased iron loading in peripheral tissues such as the liver, these *Hfe*^*-/-*^ mice exhibited no obvious iron elevation in the brain [36], raising the possibility that other factors might contribute to movement impairment with hemochromatosis-related mutations. One possibility is suggested by our previous reports of transcriptomic changes in the brain of mice with disruption of *Hfe* genes that may correlate with changes in molecular mechanisms involved in neurotransmission, synaptic plasticity, long-term potentiation and a range of other relevant systems that may affect motor control [37,38]. Another possibility, discussed further below, may be that hemochromatosis-related mutations lead to low neuronal iron despite high total brain iron.

In accordance with our previously reported quantitative [12] and histochemical [11] analyses of the *Hfe*^*-/-*^*xTfr2*^*mut*^ model, staining of ferric iron using DAB+Perls’ stain was consistent with increased iron throughout the brain, including the SNc. It is unclear if excess brain iron is a driver of PD-related neurodegeneration or instead a secondary epiphenomenon. Although high iron loading has been observed in brain regions that manifest PD-related neuropathology e.g. basal ganglia, with some MRI studies reporting cross-sectional associations between brain iron accumulation and PD, reviewed [39-42], a recent clinical trial of the brain penetrating iron chelator deferiprone accelerated, rather than delayed, PD progression [10]. The effects of iron chelators on brain and neuronal iron levels in people with hemochromatosis is unknown.

Some animal exposure models using iron supplementation by gavage or injection can achieve detectable elevation of brain iron levels, especially in the neonatal period where the blood brain barrier (BBB) is not fully developed and can also cause neurodegenerative changes relevant to PD. This has been interpreted as evidence that brain iron elevation may contribute to PD related neuropathology and motor deficits. Yet the acute, short-term nature and often high doses of iron loading in such models is not representative of brain iron loading in most human patients, where daily iron intake is far lower and tissue iron loading a slow and gradual process over extended periods [43,44]. Although using genetically-modified animals avoids the acute nature of iron loading through supplementation or injection, as noted above, many PD genetic mouse models represent extremely rare genetic conditions of dysregulated iron handling and metabolism that are unlikely to accurately model the patterns of brain iron loading seen in most people with PD.

Hemochromatosis-related mutations are relatively common. These mutations disrupt the hepcidin-ferroportin axis. The circulating peptide hepcidin regulates iron export from gut enterocytes and various other cells, including neurons, by controlling internalization and degradation of ferroportin (FPN), the only known cell surface iron exporter (reviewed [45]). High blood iron levels usually trigger hepcidin up-regulation, increasing internalization and degradation of FPN and reducing iron export from the gut into the blood, maintaining body iron balance [45]. In hemochromatosis, the normal up-regulation of hepcidin in response to high body iron loading is compromised and transport of iron by FPN from enterocytes into the blood is not appropriately down-regulated [45]. The findings in our model reported here and previously [11,12] and data from the UK BioBank [3] provide evidence that transport of iron from the blood into the brain is also increased. Determining the effects of hepcidin-ferroportin dysregulation on the human brain is challenging, as there is an ethical imperative to offer iron removal therapy (e.g. phlebotomy, chelation) as soon as patients are diagnosed. This makes it difficult to assess roles of elevated brain iron in driving PD in longitudinal assessments of community or clinical cohorts or conduct adequately-powered human studies of how iron overload conditions can increase PD susceptibility.

Powerful animal models are essential for understanding the roles of iron in PD. Surprisingly, although total brain iron is elevated in the *Hfe*^*-/-*^*xTfr2*^*mu*t^ mouse model, neurons typically remain relatively low in iron and, we speculate, might even become functionally iron deficient. In *Hfe*^*-/-*^*xTfr2*^*mu*t^ mice, liver [46] and brain hepcidin mRNA transcripts (our unpublished data) are barely detectable, suggesting brain levels of hepcidin peptide from any source, local or systemic, are likely to be negligible. We therefore anticipate minimal hepcidin-facilitated degradation of ferroportin in these mice, with maximal retention of cell surface ferroportin and consequently high iron export out of neurons and other ferroportin-expressing cells. Our published data [12] support this hypothesis – despite *Hfe*^*-/-*^*xTfr2*^*mu*t^ mice having higher total brain iron levels, suggesting iron entry into the brain is dysregulated, probably due to hepcidin deficiency, the extra iron is mainly restricted to storage reservoirs in myelinated structures and non-neuronal cells, with little histologically detected iron in neurons. We speculate that increased iron influx into the brain due to hepcidin-ferroportin dysregulation may be balanced, at least in early-stage disease, by increased iron efflux out of neurons via ferroportin, helping protect against neurotoxicity. However, over the longer term, this may lead neurons to develop functional iron deficiency.

High iron loading in hemochromatosis is usually managed by venesection or chelation. However, as noted, the chelator deferiprone accelerated PD progression in the recent PARKII trial [10]. Potential side effects in people with hemochromatosis, who may already have functional neuronal iron deficiency are not known. Determining if chelation and targeting of hepcidin affect motor control via hemochromatosis-related variants causing neuronal iron deficiency, excess or both, has direct translational value. Pre-clinical studies are needed to assess effects of chelation on brain outcomes in the context of hemochromatosis-related mutations. It is also important to test if alternative treatments targeting pathways affected by these genotypes might restore normal iron status more safely.

Finally, as noted in the Introduction, the HFE p.C282Y variant usually affects men more severely than women. This may be partly because menstruation reduces iron accumulation; risks may be higher for younger women prematurely amenorrheic due to contraceptive implants or other causes. This may lead to greater numbers of women experiencing more severe disease outcomes in the future. It will be valuable to investigate this in female mice with hemochromatosis-related mutations.

## 5 Conclusion

In addition to elevated brain iron levels, as previously reported [11,12], hemochromatosis-related mutations caused features relevant to PD in male mice, specifically depletion of SNc dopaminergic TH^+^ neurons at both 3 months and 9 months of age and pronounced deficits in balance and motor coordination by 9 months. These features place this model at the forefront of genetic mouse models with parkinsonian features. Further research is required to determine whether these changes are attributable to chronic brain iron loading alone or other effects of hemochromatosis-related mutations. Gaining a better understanding of the mechanisms underlying parkinsonian features in these mice is likely to help guide preventive or therapeutic strategies for men at increased risk of PD due to hemochromatosis-related mutations and may also provide insights into sporadic PD.

## Author Contributions

**Elvis Acquah:** Investigation, Formal analysis, Validation, Visualization, Writing - Original Draft, Review & Editing. **Rebecca Hood**: Methodology, Investigation, Formal analysis, Validation, Visualization, Writing - Original Draft, Review & Editing. **Chan-An Lin**: Software, Investigation, Visualization, Writing - Review & Editing. **Qiao-Xin Li**: Investigation, Visualization, Supervision, Writing - Review & Editing. **Luke Gordon:** Investigation, Validation. **Ritambhara Aryal:** Writing - Review & Editing. **Kristy Martin:** Investigation: Writing - Review & Editing. **Adrienne Milward**: Conceptualization, Supervision, Writing - Original Draft, Review & Editing. **Daniel Johnstone:** Conceptualization, Methodology, Project administration, Supervision, Writing - Original Draft, Review & Editing.

## Acknowledgements

This research was supported by Australian Government Research Training Program scholarships to EA, LG and RA; RH is supported by the Neurosurgical Research Foundation.

